# Comparative thermophysiology of marine *Synechococcus* CRD1 strains isolated from different thermal niches in iron-depleted areas

**DOI:** 10.1101/2022.03.09.483587

**Authors:** Mathilde Ferrieux, Louison Dufour, Hugo Doré, Morgane Ratin, Audrey Guéneuguès, Léo Chasselin, Dominique Marie, Fabienne Rigaut-Jalabert, Florence Le Gall, Théo Sciandra, Garance Monier, Mark Hoebeke, Erwan Corre, Xiaomin Xia, Hongbin Liu, David J. Scanlan, Frédéric Partensky, Laurence Garczarek

**Author notes:** Department of Ecology, Evolution and Marine Biology; University of California, Santa Barbara, USA. **Correspondence:** Corresponding Author: L. Garczarek.

## Abstract

Marine *Synechococcus* cyanobacteria are ubiquitous in the ocean, a feature likely related to their extensive genetic diversity. Amongst the major lineages, clades I and IV preferentially thrive in temperate and cold, nutrient-rich waters, whilst clades II and III prefer warm, nitrogen or phosphorus-depleted waters. The existence of such cold (I/IV) and warm (II/III) thermotypes is corroborated by physiological characterization of representative strains. A fifth clade, CRD1, was recently shown to dominate the *Synechococcus* community in iron-depleted areas of the world ocean and to encompass three distinct ecologically significant taxonomic units (ESTUs CRD1A-C) occupying different thermal niches, suggesting that distinct thermotypes could also occur within this clade.

Here, using comparative thermophysiology of strains representative of these three CRD1 ESTUs we show that the CRD1A strain MITS9220 is a warm thermotype, the CRD1B strain BIOS-U3-1 a cold temperate thermotype, and the CRD1C strain BIOS-E4-1 a warm temperate stenotherm. Curiously, the CRD1B thermotype lacks traits and/or genomic features typical of cold thermotypes. In contrast, we found specific physiological traits of the CRD1 strains compared to their clade I, II, III and IV counterparts, including a lower growth rate and photosystem II maximal quantum yield at most temperatures and a higher turnover rate of the D1 protein. Together, our data suggests that the CRD1 clade prioritizes adaptation to low-iron conditions over temperature adaptation, even though the occurrence of several CRD1 thermotypes likely explains why the CRD1 clade as a whole occupies most iron-limited waters.

## INTRODUCTION

Marine picocyanobacteria contribute to the biogeochemical cycling of various elements, most notably carbon, contributing ~25% of ocean net primary productivity, of which the *Synechococcus* genus alone is responsible for about 16% (Flombaum et al., 2013). The large geographic distribution of these organisms, extending from the equator to subpolar waters, is largely attributable to their extensive genetic and functional diversity (Zwirglmaier et al., 2008; Farrant et al., 2016; Doré et al., 2020). Amongst the nearly 20 clades within subcluster (SC) 5.1, the most abundant and diversified *Synechococcus* lineage in oceanic ecosystems (Dufresne et al., 2008; Scanlan et al., 2009; Ahlgren and Rocap, 2012), only four (clades I, II, III and IV) were thought to largely dominate *in situ*. Clades I and IV mainly thrive in temperate and cold, nutrient-rich waters, while clades II and III reside in warm, oligotrophic or mesotrophic areas (Zwirglmaier et al., 2008; Mella-Flores et al., 2011), suggesting the existence of cold (I/IV) and warm (II/III) *Synechococcus* ‘thermotypes’. This hypothesis was subsequently confirmed by work demonstrating that strains representative of these different clades exhibit distinct thermal *preferenda* (Mackey et al., 2013; Pittera et al., 2014; Breton et al., 2020; Six et al., 2021), a feature notably linked to differences in the thermostability of light-harvesting complexes (Pittera et al., 2017), lipid desaturase gene content (Pittera et al., 2018) and the ability of some strains to induce photoprotective light dissipation at colder temperatures using the orange carotenoid protein (OCP; Six et al., 2021). Field studies using global ocean datasets have allowed to refine the respective ecological niches of the different thermotypes, with clade I extending further north than clade IV (Paulsen et al., 2016; Doré et al., 2022) and clades II and III predominating in N- and P-depleted waters, respectively, but also to highlight the importance of a fifth clade within SC 5.1, the CRD1 clade (Farrant et al., 2016; Sohm et al., 2016; Kent et al., 2019). Initially thought to be limited to the Costa Rica dome area (Saito et al., 2005; Gutiérrez-Rodríguez et al., 2014), the latter clade was recently found to be a major component of *Synechococcus* communities in Fe-depleted areas (Farrant et al., 2016; Sohm et al., 2016; Ahlgren et al., 2020). Furthermore, analysis of the global distribution of these organisms using high-resolution marker genes has highlighted large within-clade microdiversity associated with niche differentiation in marine *Synechococcus* (Farrant et al., 2016; Larkin and Martiny, 2017; Xia et al., 2019), as also observed in *Prochlorococcus* (Kashtan et al., 2014; Larkin et al., 2016). Using the *petB* gene encoding cytochrome *b*_6_, Farrant et al. (2016) showed that most major clades encompassed several Ecologically Significant Taxonomic Units (ESTUs), i.e. genetically related subgroups within clades occupying distinct oceanic niches. This is notably the case for ESTU IIB that occupies a cold thermal niche in sharp contrast with IIA, the dominant ESTU within clade II that occupies warm, mesotrophic, and oligotrophic iron (Fe)-replete waters. Similarly, three distinct ESTUs with distinct thermal niches were identified within the CRD1 clade and the co-occurring clade EnvB (a.k.a. CRD2; Ahlgren et al., 2020): i) CRD1B/EnvBB are found in cold mixed waters in co-occurrence with ESTUs IA, IVA and IVC, ii) CRD1C/EnvBC dominate in warm, high-nutrient low-chlorophyll (HNLC) regions such as the central Pacific Ocean, and iii) CRD1A/EnvBA are present in both environments and thus span a much wider range of temperatures than CRD1B and C (Farrant et al., 2016). This suggests that these three CRD1 ESTUs could correspond to different thermotypes.

In order to test this hypothesis, we used strains representative of each of the three CRD1 ESTUs to determine the fundamental thermal niches of these organisms as compared to typical cold (clades I and IV) and warm (clades II and III) thermotypes. Furthermore, given the strong influence of temperature on optimal functioning of the photosynthetic apparatus in marine *Synechococcus* (Pittera et al., 2014, 2017; Guyet et al., 2020), we also examined the effect of temperature acclimation on the photophysiology of CRD1 ESTUs compared to their clade I and IV counterparts and show that CRD1 thermotypes actually differ more strongly in this respect to members of clades I-IV than from each other.

## MATERIALS AND METHODS

### Strains and growth conditions

The eight *Synechococcus* spp. strains used in this study were retrieved from the Roscoff Culture Collection (RCC; https://roscoff-culture-collection.org/), including representative strains of the three known CRD1 ESTUs (CRD1A – C) and one or two of each of the four dominant clades in the field (clades I – IV) used as controls (Table 1 and Supplementary Fig. 1). Cells were grown in 50 mL flasks (Sarstedt, Germany) in PCR-S11 culture medium (Rippka et al., 2000) supplemented with 1 mM sodium nitrate. Cultures were acclimated in temperature-controlled chambers across a range of temperatures dependent on the thermal tolerance of each strain. Continuous light of 75 μmol photons m^−2^ s^−1^ (hereafter μE m^−2^ s^−1^) was provided by a white-blue-green LED system (Alpheus, France). For each experiment cultures were grown in triplicate, inoculated at an initial cell density of ~3 x 10^6^ cells mL^−1^, and samples harvested daily to measure growth rate and fluorescence parameters as described below.

**TABLE 1.** Characteristics of the *Synechococcus* strains used in this study

| Strains name | MVIR-18-1 | A15-62 | M16.1 | WH8102 | BL107 | BIOS-E4-1 | BIOS-U3-1 | MITS9220 |
| --- | --- | --- | --- | --- | --- | --- | --- | --- |
| RCC # <sup>1</sup> | 2385 | 2374 | 791 | 539 | 515 | 2534 | 2533 | 2571 |
| Subcluster <sup>2</sup> | 5.1 | 5.1 | 5.1 | 5.1 | 5.1 | 5.1 | 5.1 | 5.1 |
| Clade <sup>2</sup> | I | II | II | III | IV | CRD1 | CRD1 | CRD1 |
| ESTU <sup>2</sup> | IB | IIA | IIA | IIIA | IVA | CRD1C | CRD1B | CRD1A |
| Pigment type <sup>3</sup> | 3aA | 3dB | 3a | 3c | 3dA | 3cA | 3dA | 3dA |
| Ocean | Atlantic | Atlantic | Atlantic | Atlantic | Med. Sea | Pacific | Pacific | Pacific |
| Region | North Sea | Offshore Mauritania | Gulf of Mexico | Caribbean Sea | Balearic Sea | South East Pacific | Chile upwelling | Equatorial Pacific |
| Isolation latitude | 61°00' N | 17°37' N | 27°70' N | 22°48' N | 41°72' N | 31°52' S | 34°00' S | 0°00' N |
| Isolation longitude | 1°59' E | 20°57' W | 91°30' W | 65°36' W | 3°33' E | 91°25' W | 73°22' W | 140°00' W |

<sup>1</sup>Roscoff Culture Collection, <sup>2</sup>Farrant et al. (2016), <sup>3</sup>Humily et al. (2013).

In order to compare the capacity of strains to repair the D1 subunit of photosystem II (PSII; see ‘Measurement of PSII repair rate’ section), cultures grown in 250 ml flasks were acclimated at 18, 22 and 25°C, temperatures at which all strains were able to grow, were subjected to high light stress (350 μE m^−2^ s^−1^). Exponentially growing cultures were sampled at T0 and after 15, 30, 60, and 90 min of stress, before shifting cultures back to the initial light conditions and then sampling again after 15, 30, 60 min and 24h of recovery (R). While D1 repair measurements were performed at all time points, cell concentrations were measured by flow cytometry only at T0, T30min, T90min, R30min and R24h and liposoluble pigment content was determined only at T0.

### Flow cytometry

Culture aliquots (200 μl) sampled for flow cytometry were fixed using 0.25% (v/v) glutaraldehyde (grade II, Sigma Aldrich, USA) and stored at −80°C until analysis (Marie et al., 1999). Cell concentrations were estimated using a Guava easyCyte flow cytometer (Luminex Corporation, USA) and maximum growth rates (μ_max_) were calculated as the slope of the linear regression of ln(cell density) *vs*. time during the exponential growth phase. *Synechococcus* cells were identified based on their red (695 nm) and orange (583 nm) fluorescence, proxies for their chlorophyll *a* and phycoerythrin content, respectively. Fluorescence, forward scatter and side scatter values were normalized to that of standard 0.95 μm beads using Guavasoft software (Luminex Corporation, USA).

### Fluorescence measurements

The maximum PSII quantum yield (F_v_/F_M_) was estimated using a Pulse Amplitude Modulation fluorimeter (Phyto-PAM II, Walz, Germany) during the exponential growth phase after adding 100 μM of the PSII blocker 3-(3,4-dichlorophenyl)-1,1-dimethylurea (DCMU, Sigma-Aldrich, USA).

The PSII quantum yield was calculated as:

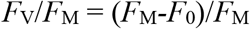

where *F*_0_ is basal fluorescence, *F*_M_ maximal fluorescence level and *F*_V_ variable fluorescence (Campbell et al., 1998; Six et al., 2007).

Fluorescence excitation (with emission set at 580 nm) and emission (with excitation set at 530 nm) spectra were generated using a LS-50B spectrofluorometer (Perkin-Elmer, USA) as described in Six et al. (2004). The fluorescence excitation ratio (Exc_495:550nm_) was used as a proxy for the PUB:PEB ratio. Phycobilisome (PBS) rod length and the degree of coupling of the PBS to PSII reaction center chlorophylls was then assessed using fluorescence emission spectra by calculating the phycoerythrin (PE, *F*_max_ = 565–575 nm) to phycocyanin (PC, *F*_max_ = 645–655 nm) ratio as well as the PC to PBS terminal acceptor (TA; *F*_max_ = 680 nm) ratio, respectively (Pittera et al., 2017).

### Pigment analyses

Triplicate cultures were harvested during the exponential phase when *F*_v_/*F*_M_ was maximum for each temperature condition. Cultures (50 mL) were subjected to centrifugation in the presence of 0.01% (v/v) pluronic acid (Sigma-Aldrich, Germany) at 4°C, 14,000 x *g* for 7 min, using an Eppendorf 5804R (Eppendorf, France). Pellets were resuspended and transferred to 1.5 ml Eppendorf tubes and centrifuged at 4°C, 17,000 x g for 2 min using an Eppendorf 5417R centrifuge (Eppendorf, France). Once the supernatant was removed samples were stored at −80 °C until further analysis. Pigment content was subsequently assessed using calibrated high-performance liquid chromatography (HPLC 1100 Series System, Hewlett Packard, St Palo Alto, CA), as previously described (Six et al., 2005).

### Measurement of the photosystem II repair rate

Each culture acclimated to 75 μE m^−2^ s^−1^ and 18, 22 or 25 °C was split into two new 50 mL flasks (Sarstedt Germany) with one used as a control and the other flask supplemented with the ribosome blocker lincomycin (0.5 mg mL^−1^ final concentration, Sigma-Aldrich, USA). Both sub-cultures were then subjected to light stress by exposing cultures to 375 μE m^−2^ s^−1^ continuous light (at the same temperature), and *F*_V_/*F*_M_ measured at different time points as described above. The PSII repair rate for each strain at each temperature was determined from the coefficient differences between the exponential curves fitted over the 90 min time course of *F*_V_/*F*_M_ measurements for control and +lincomycin samples. Each experiment was done in quadruplicate.

### Determination of the realized environmental niches of major *Synechococcus* ESTUs

The realized niches of CRD1 and clades I-IV ESTUs were determined using *petB* reads extracted from metagenomic data from the *Tara* Oceans and *Tara* Polar circle expeditions, the Ocean Sampling Day (OSD; June 21^st^ 2014) campaign, and *petB* metabarcodes from i) various oceanographic cruises (CEFAS, BOUM, Micropolar, RRS Discovery cruise 368 and several in the northwestern Pacific Ocean as detailed in Xia et al., 2017), ii) two individual sampling sites in the Mediterranean Sea (Boussole, Point B) as well as iii) a bi-monthly sampling at the long-term observatory site SOMLIT (“Service d’Observation en Milieu Littoral”)-Astan located 2.8 miles off Roscoff between July 2009 and December 2011 (Supplementary Table 1).

*petB* metagenomic recruitment using the *Tara* Oceans and OSD datasets was performed as described previously (Farrant et al., 2016). *Synechococcus petB* sequences from both metagenomes and metabarcodes were used to define operational taxonomic units (OTUs) at 97% identity using Mothur v1.34.4 (Schloss et al., 2009) that were then taxonomically assigned using a *petB* reference database (Farrant et al., 2016). OTUs encompassing more than 3% of the total *Synechococcus* reads for a given sample were grouped into ESTUs and used to determine the whole temperature range occupied by each of the five major *Synechococcus* ESTUs.

### Comparative genomics

The Cyanorak *v2.1* information system (http://www.sb-roscoff.fr/cyanorak/; Garczarek et al., 2021) was used to compare the phyletic pattern i.e., the presence/absence pattern of each cluster of likely orthologous genes (CLOG) in each strain, for CRD1 strains and their clades I-IV counterparts for a number of selected genes potentially involved in adaptation to low temperature.

## RESULTS

### The fundamental thermal niches of CRD1 *vs*. clades I to IV strains

In order to determine the temperature optima and boundary limits of the different CRD1 strains and to compare them to those of typical cold and warm *Synechococcus* thermotypes, representative strains of each of the three CRD1 ESTUs and strains of clades I, II, III and IV were grown over a range of temperatures from 6 to 36°C. The growth responses of all strains to temperature followed a typical asymmetric bell-shaped curve over the selected temperature range (Fig. 1), with a progressive rise in growth rate (μ) until *T*_opt_ (the temperature associated with maximum μ: μ_max_) was reached, and a sharp decline above *T*_opt_. BIOS-U3-1 (CRD1-B) was able to grow between 12 and 29°C with a *T*_opt_ at 25°C (μ_max_ = 0.78 ± 0.02 d^−1^), a growth pattern most similar to that of the clade IV strain BL107, while the clade I strain MVIR-18-1 was able to grow at much lower temperatures, down to 8°C but could not grow above 25°C (Fig. 1A). MITS9220 (CRD1-A) and BIOS-E4-1 (CRD1-C) displayed thermal growth characteristics more similar to the clade II (A15-62 and M16.1) and III (WH8102) strains, representatives of warm thermotypes (Fig. 1B). While most strains in this category displayed a minimal growth temperature of 16°C, large variations between strains were observed at the highest thermal boundary limit (*T*_max_). Maximum growth temperature was obtained for M16.1 (II; *T*_max_: 34°C), then A15-62 (II) and WH8102 (III; both with *T*_max_ at 32°C), MITS9220 (CRD1-A; *T*_max_: 31°C) and finally for BIOS-E4-1 (CRD1-C; *T*_max_: 30°C). The latter strain also displayed the highest minimal growth temperature (*T*_min_: 18°C) and thus possesses the narrowest temperature range for growth of all the strains studied (12°C *vs*. 15-18°C). It is also worth noting that CRD1 strains display a lower maximum growth rate and more generally lower growth rates at most temperatures than their clade I, II, III and IV counterparts.

**FIGURE 1.**
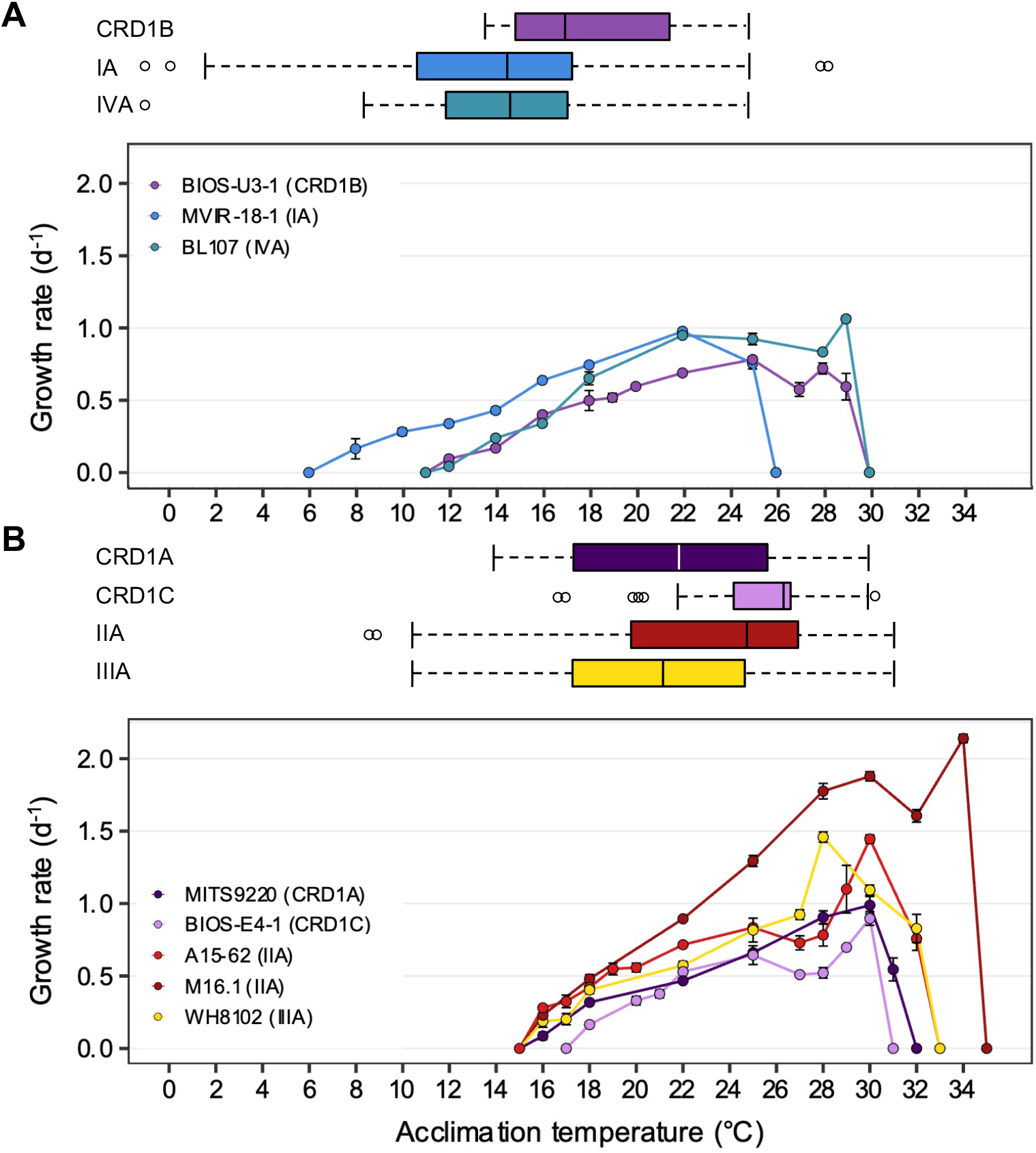
Fundamental thermal niches of CRD1 *vs*. clade I, II, III and IV strains and environmental realized niches of the corresponding ESTUs. **(A)** CRD1-B strain BIOS-U3-1 *vs*. cold thermotypes. **(B)** CRD1-A strain MITS9220 and the CRD1-C strain BIOS-E4-1 *vs*. warm thermotypes. The insert indicates the strain names and their corresponding ESTU (*sensu* Farrant et al., 2016) between brackets. Each data point is the average of at least 3 biological replicates. Environmental realized niches are indicated as horizontal boxplots for each ESTU above each graph with the whiskers corresponding to 1.5-fold the interquartile range and outliers being plotted as individual points.

### The environmental realized niches of CRD1 *vs*. clades I to IV strains

We then compared the fundamental thermal niches of all studied strains, i.e. the whole temperature range over which they can grow in a laboratory setting in the absence of biotic interactions (e.g. competition or predation), with environmental realized niches (*sensu* Pearman et al., 2008) of the corresponding ESTUs. For this, we determined the distribution limits of each of these ESTUs along the *Tara* Oceans and *Tara* Polar circle transects, 203 samples from OSD2014 and additional oceanographic cruises and individual sampling sites, altogether encompassing 421 samples worldwide covering a wide range of temperature conditions (Fig. 1, Supplementary Fig. 2, Supplementary Table 1). This made it possible to have much finer estimates of the limits of the thermal niches of the different ESTUs than in the study performed by Farrant et al. (2016), in particular for the cold adapted ESTUs, which were poorly represented in the initial *Tara* Oceans dataset (Supplementary Fig. 2).

This analysis showed that the CRD1B ESTU displayed a reduced thermal tolerance range in the environment (14 to 24.5°C) compared to the BIOS-U3-1 strain in culture (12 to 29°C), while the typical cold thermotypes colonized larger thermal niches *in situ* than their representative strains (Fig. 1A). Environmental realized niches indeed ranged from 2.5 to 24°C for ESTU IA (compared to 8 to 25°C for MVIR-18-1) and from 8.5 to 25°C for ESTU IVA (compared to 12 to 29°C for BL107). Interestingly, the median temperature of the CRD1B ESTU is 3°C higher than that observed for ESTUs IA and IVA.

As concerns warm thermotypes, CRD1C displayed a fairly narrow thermal tolerance range *in situ* (22 to 29.5°C), which, similar to the cold thermotype CRD1B, was even narrower than for its representative strain BIOS-E4-1 (18 to 30°C; Fig. 1B). Comparatively, the CRD1A ESTU was detected across a wider temperature range (14 to 30.5°C) than the other two CRD1 ESTUs and also slightly larger than the corresponding strain in culture (MITS9220, 16 to 31°C). Still, the most extended temperature range was observed for ESTU IIA and IIIA (12 to 32°C) that reached significantly lower temperature limits than the corresponding clade II (16 to 32-34°C) and III (16 to 32°C) strains. Of note, although both IIA and IIIA ESTUs displayed a similar temperature range, the median temperature of ESTU IIA (25°C) was about 3°C higher than that of ESTU IIIA (22°C) and the maximum median temperature was surprisingly observed for the CRD1C ESTU (26.5°C). In this context, it is also worth mentioning that although clade II strains are clearly both warm thermotypes, M16.1 displays a significantly higher temperature limit for growth than A15-62 and more generally than all other strains. This suggests that ESTU IIA may encompass two distinct ESTUs, but such a high temperature niche (>32 °C) where they could be discriminated is exceptional and not available in our dataset (Supplementary Table 1).

### Comparative genomics

In order to assess whether the cold, temperate thermotype BIOS-U3-1 (CRD1B) exhibits similar adaptation mechanisms to those previously described for the cold-adapted clades I and/or IV, we examined a number of clusters of likely orthologous genes (CLOGs) from all *Synechococcus* genomes belonging to clades I-IV and CRD1 present in the Cyanorak *v2.1* information system (Garczarek et al., 2021). First, we looked for the occurrence of two amino-acid substitutions in phycocyanin α- (RpcA) and ß-subunits (RpcB), which were shown to differ between cold- (Gly in clades I and IV for RpcA43; Ser in RpcB42) and warm-thermotypes (Ala in clades II and III for RpcA43; Asp in RpcB42), these substitutions being potentially responsible for the differential thermotolerance of this phycobiliprotein between thermotypes (Pittera et al., 2017). In all three CRD1 strains, both sites displayed the warm-type residue (Supplementary Fig. 3A-B), suggesting that in contrast to typical cold and warm thermotypes, the molecular flexibility of this phycobiliprotein does not differ between CRD1 thermotypes. We then looked at fatty acid desaturases that are essential for regulating membrane fluidity and thus the activity of integral membrane proteins, including photosynthetic complexes (Mikami and Murata, 2003; Pittera et al., 2018; Breton et al., 2020). All three CRD1 strains surprisingly possess in addition to the core Δ9-desaturase gene *desC3,* a second Δ9-desaturase, *desC4* (Supplementary Table 2), previously thought to be specific to cold-adapted strains as well as the Δ12-desaturase *desA3* found in both cold-adapted clades I and IV as well as in clade III, a warm thermotype subjected to much stronger seasonal variability than its (sub)tropical clade II counterparts (Pittera et al., 2018). Furthermore, BIOS-U3-1 also possesses *desA2,* thought to be specific to warm environments, while this gene is in contrast absent from the other two CRD1 warm-adapted strains. Thus, CRD1 strains exhibit a different desaturase gene set and potentially display a larger capacity to regulate membrane fluidity than typical cold- or warm-adapted thermotypes. Finally, while all clades I, III and IV genomes possess the *ocp* operon, involved in the protection of PSII against photoinactivation through the dissipation of excess light energy (Kirilovsky, 2007) and which was recently shown in marine *Synechococcus* to play a key role at low temperature (Six et al., 2021), none of the three CRD1 genomes possess this operon.

### Photosynthetic activity and pigment content

PSII quantum yield (F_V_/F_M_), used as a proxy of photosynthetic activity, was measured for each strain over their whole temperature growth range. Most strains displayed a decrease in this parameter at both low and high boundary limits of their growth temperature range and this effect was particularly striking for BIOS-U3-1, reaching values down to 0.11 at 14°C and 0.32 at 28°C (Fig. 2A). Besides MVIR-18-1 that exhibited a quite constant F_V_/F_M_ over its whole temperature range, the decrease in F_V_/F_M_ at high temperature was stronger for cold than warm thermotypes that are able to maintain a quite high F_V_/F_M_ in the warmest growth conditions (Fig. 2B). Finally, as for growth rate, CRD1 strains exhibited lower F_V_/F_M_ at all temperatures than clade I to IV strains.

**FIGURE 2.**
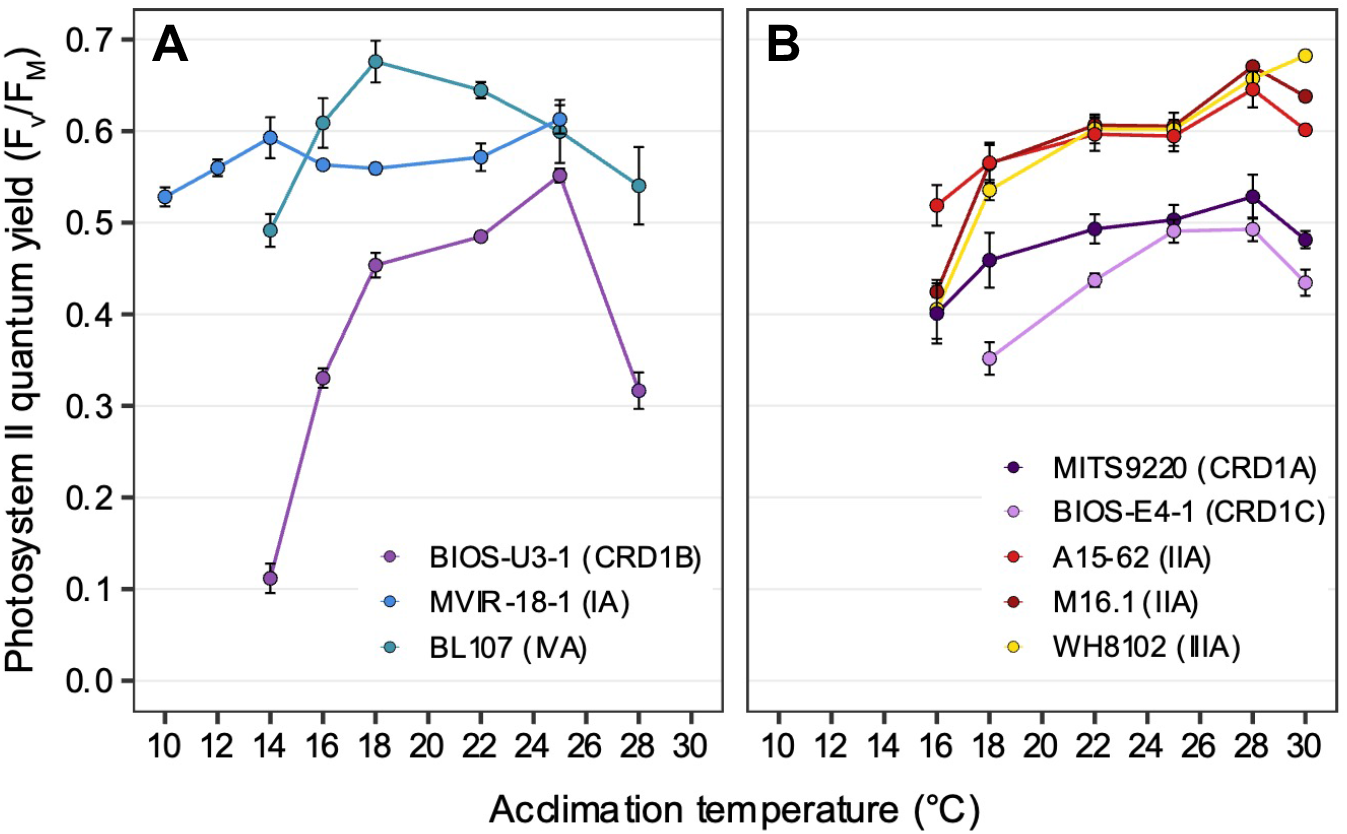
Variation with growth temperature of photosystem II quantum yield (F_V_/F_M_) for CRD1 *vs*. clade I, II, III and IV strains. **(A)** CRD1-B strain BIOS-U3-1 *vs*. cold thermotypes. **(B)** CRD1-A strain MITS9220 and the CRD1-C strain BIOS-E4-1 *vs*. warm thermotypes. The insert indicates the strain names and their corresponding ESTU (*sensu* Farrant et al., 2016) between brackets.

The Exc_495:550nm_ fluorescence excitation ratio, used as a proxy for PUB:PEB ratio, was consistent with the pigment type of each strain (Humily et al., 2013; Table 1; Supplementary Fig. 4A). This ratio remained pretty constant over the whole temperature range for all strains except for the chromatic acclimator BL107 (pigment type 3dA), for which a sharp increase was observed at its maximal growth temperature (28°C) to reach a value (1.35) intermediate between that typically observed in green light (or white light; 0.6-0.7) and blue light (1.6-1.7). This suggests that the chromatic acclimation process could be affected by growth temperature, at least in this strain. The phycobilisome (PBS) rod lengths and the degree of coupling of PBS to PSII reaction center chlorophylls, as estimated from PE:PC and PE:TA ratios respectively, showed fairly limited variations over the temperature range, indicating that the phycobiliprotein composition of PBS is quite stable over the growth temperature range of each strain (Supplementary Fig. 4B-C). One notable exception was a rise in both ratios for strain A15-62 at its minimal growth temperature, likely attributable to the partial decoupling of individual phycobiliproteins and of the whole PBS from PSII, a phenomenon typically observed under stressful conditions (Six et al., 2007; Guyet et al., 2020). It is also worth noting that MITS9220 and to some extent BIOS-E4-1, exhibited a significantly higher PE:PC ratio than the five other strains, potentially indicating a different phycobiliprotein composition and/or length of PBS rods.

In terms of liposoluble pigments, the ß-carotene/chlorophyll *a* (β-car/Chl *a*) ratio tended to increase with temperature in BIOS-E4-1 and MITS9220, as observed for the other warm thermotypes, whilst this ratio was more stable in the cold thermotypes BIOS-U3-1 and BL107, and seemed to slightly increase in the lower part of the thermal range for the clade I strain MVIR-18-1 (Fig. 3). For all strains, these ratios result from a concomitant increase with temperature of Chl *a* and β-car content per cell (Supplementary Fig. 5), indicating an enhancement of the surface of thylakoids per cell at higher temperatures that was particularly marked for BIOS-E4-1 and A15-62, whilst this variation was fairly limited in the other two CRD1 strains. As these two pigments are present in different proportions in PSI and II (Umena et al., 2011; Xu and Wang, 2017), the higher β-car/Chl a ratio measured in clades I and IV strains also suggests that they may have a higher PSII:PSI ratio than all other strains, including BIOS-U3-1, and that this ratio might be more strongly affected by temperature in warm than cold thermotypes.

**FIGURE 3.**
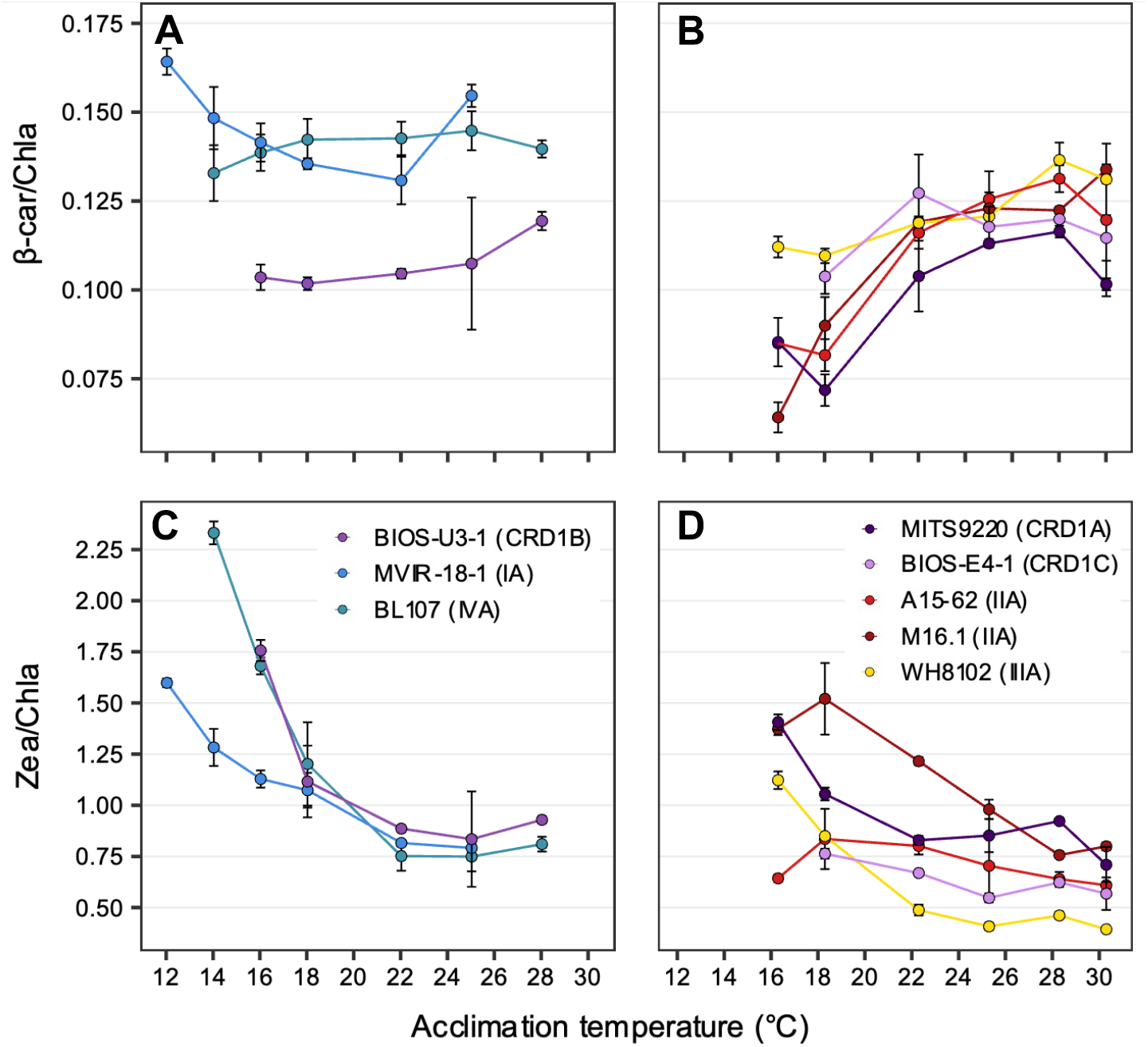
Variation with growth temperature of cellular mass pigment ratios of CRD1 *vs*. clade I, II, III and IV strains. **(A, B)** β-carotene (β-car) to chlorophyll *a* (Chla) ratio. **(C, D)** Zeaxanthin (Zea) to Chl*a* ratio with **(A, C)** CRD1-B strain BIOS-U3-1 *vs*. cold thermotypes and **(B, D)** CRD1-A strain MITS9220 and the CRD1-C strain BIOS-E4-1 *vs*. warm thermotypes. Inserts indicate strain names and their corresponding ESTUs (*sensu* Farrant et al., 2016) between brackets.

As concerns the zeaxanthin/chlorophyll *a* (Zea/Chl *a*) ratio, although an increase in this ratio was measured at low temperature for all strains, the amplitude was globally larger for cold than for warm thermotypes, with BIOS-U3-1 behaving very similarly to the clade IV strain BL107 that exhibits the largest variation in this ratio (Fig. 3). Changes in this ratio likely originate partially from the decrease in Chl *a* content in response to cold, a strategy typically used by cells to regulate light utilization under slow growth conditions (Inoue et al., 2001). However, several strains also displayed an increase in their Zea content per cell at low temperature, a response particularly striking in BIOS-U3-1 and A15-62, but that also seems to occur in M16.1 and in the two other CRD1 strains BIOS-E-4-1 and MITS9220 (Supplementary Fig. 5). Thus, although Zea has been hypothesized to be involved in the photoprotection of cold-adapted strains by dissipating excess light energy under low temperature conditions (Kana et al., 1988; Breton et al., 2020), this process seems to be present in both cold and warm-adapted CRD1 strains and in most warm thermotypes as well. In this context, it is also worth noting that the two clade II strains, A15-62 and M16.1, displayed fairly distinct temperature-induced variations in their Zea:Chl *a* ratios and individual pigment contents, possibly linked to their different isolation temperatures (see discussion below).

### Photosystem II repair capacity

The ability of the different strains to repair PSII in response to light stress (375 μE m^−2^ s^−1^) was determined in cultures acclimated to 18, 22 and 25°C by measuring changes in F_V_:F_M_ over time after adding the protein synthesis inhibitor lincomycin, or not (Supplementary Fig. 6). While a decrease in F_V_:F_M_ ratio during the 90 min light stress period was observed in both cultures supplemented with lincomycin and controls, this ratio only re-increased back up to initial F_V_:F_M_ values, after shifting cultures back to standard light conditions (75 μE m^−2^ s^−1^), in the control group in most strains and temperature conditions. Thus, all studied strains were able to recover from this light stress, as long as the D1 repair cycle was not inactivated by inhibition of protein synthesis. Yet, a fast decrease in F_V_:F_M_ was observed for all three CRD1 cultures supplemented with lincomycin, while the +/− lincomycin curves overlapped during the first 15-30 min of light stress in most other strains and conditions. This suggests that the initial decrease in F_V_:F_M_ in clades I-IV strains was not due to D1 damage but rather to dissipation of light energy as heat through non-photochemical quenching (Campbell et al., 1998), whilst the damage and hence repair of D1 proteins only occurred later on.

The PSII repair rate (R_PSII_), as calculated from the time course of F_V_:F_M_ with and without lincomycin, increased with temperature in most strains, except for BIOS-U3-1 that displayed its highest rate at 22°C (Fig. 4). Strikingly, all three CRD1 strains displayed significantly higher R_PSII_ than clade I-IV strains at all three tested temperatures, a difference ranging from 3-to nearly 40-fold at the lowest common temperature (18°C). Furthermore, CRD1 strains displayed fairly limited variation in R_PSII_ with temperature (ranging from 1.33 to 1.87-fold) compared to the other strains, the strongest increase in R_PSII_ being observed for the clade I strain MVIR-18-1 (21.5-fold) and the clade III strain WH8102 (5.5-fold). This indicates that CRD1 strains exhibit a constitutively high level of PSII repair compared to the other strains whatever the growth temperature and only trigger a moderate increase in R_PSII_ in response to temperature variations.

**FIGURE 4.**
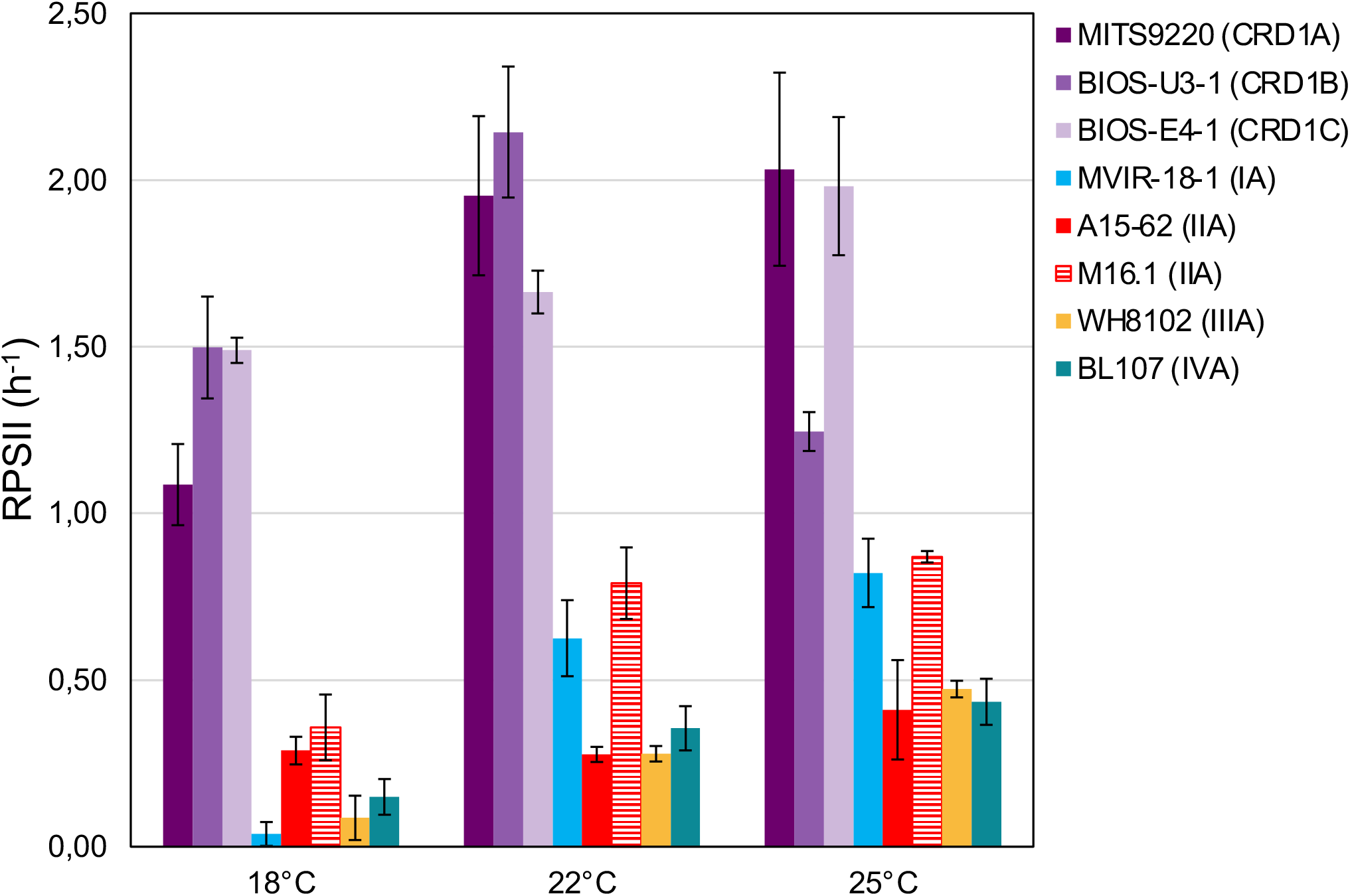
Variation with growth temperature of cumulative photosystem II repair rate (RPSII) of CRD1 *vs*. clade I, II, III and IV strains. The insert indicates the strain names and their corresponding ESTU (*sensu* Farrant et al., 2016) between brackets.

## DISCUSSION

Temperature constitutes one the strongest driving factors that have shaped genetic diversification and niche partitioning in marine cyanobacteria (Scanlan et al., 2009; Flombaum et al., 2013; Biller et al., 2015) and phytoplankton at large (Sunagawa et al., 2015; Delmont et al., 2020). While temperature has caused one major diversification event in *Prochlorococcus,* resulting in the divergence of the cold-adapted HLI from the warm-adapted HLII clades (Johnson et al., 2006; Kettler et al., 2007), several independent temperature-related diversification events also occurred in the *Synechococcus* SC 5.1 radiation, leading to the emergence of clades I and IV (Dufresne et al., 2008; Zwirglmaier et al., 2008). Here, determination of the temperature optimum and boundary limits (i.e., the fundamental niche) of strains representative of the three CRD1 ESTUs identified in the field (Farrant et al., 2016) showed that different thermotypes can also be delineated within the CRD1 clade, which dominates the *Synechococcus* populations in low-Fe areas of the world Ocean. Comparison with representative strains of the cold-adapted *Synechococcus* ecotypes (clades I and IV) on the one hand, and warm-adapted ecotypes (clades II and III) on the other, made it possible to classify i) the CRD1A strain MITS9220, isolated from the equatorial Pacific Ocean, as a warm thermotype, ii) the CRD1B strain BIOS-U3-1, isolated from the Chilean upwelling, as a cold temperate thermotype, and finally iii) the CRD1C strain BIOS-E4-1, isolated from the edge of the South Pacific gyre, a stable, warm, Fe-depleted oceanic region (Claustre et al., 2008), as a warm temperate stenotherm.

As expected from theory (Pearman et al., 2008), the realized environmental thermal niches of CRD1 ESTUs were narrower than their fundamental niches (or similar for MITS9220). In contrast, for ESTUs IA to IVA, the realized environmental niche was significantly more extended towards the low thermal limit than the fundamental niches of their representative strains, this expansion being particularly marked for ESTU IA (Fig. 1). This could be due to passive transport of *Synechococcus* populations by currents into water masses colder than their temperature limits for growth. Alternatively, these ESTUs may exhibit a greater microdiversity than previously assessed (Farrant et al., 2016) and could be subdivided into distinct ESTUs occupying slightly different thermal niches from the current ones, although representative strains to test this hypothesis remain to be isolated. In agreement with the latter hypothesis Paulsen et al. (2016) measured a positive growth rate of *Synechococcus* natural populations dominated by clade I in waters as cold as 2°C in the vicinity of the Svalbard island. Thus, CRD1 ESTUs appear to be strongly outcompeted by their ESTU IA to IVA counterparts at their lower temperature limits. Consistent with this, comparison of their gene content showed that CRD1 ESTUs, including the cold thermotype CRD1B, lack the main adaptation mechanisms reported so far for typical cold thermotypes. Indeed, all CRD1 strains examined in this study i) exhibit warm-type substitutions in their α and β-phycocyanin subunits, influencing the thermotolerance of this phycobiliprotein (Pittera et al., 2017; Supplementary Fig. 3A-B); ii) possess a different set of desaturase genes, involved in regulation of membrane fluidity (Pittera et al., 2018), than typical warm and cold thermotypes (Supplementary Table 2) and iii) lack the OCP system, involved in the protection of PSII against photoinactivation, which seemingly plays a key role at low temperature (Kirilovsky, 2007; Six et al., 2021). Still, we cannot exclude that CRD1 strains could use alternative strategies to cope with temperature variations and notably to deal with the generation of reactive oxygen species, known to be generated by a variety of factors including low and high temperature (Nishiyama et al., 2006; Latifi et al., 2009). For instance, all CRD1 strains possess the *srxA* gene encoding sulfiredoxin catalyzing the reduction of 2-Cys peroxiredoxin involved in H_2_O_2_ detoxification (Findlay et al., 2005; Guyet et al., 2020) as well as *isiA* that, besides its role in increasing the light-harvesting efficiency of PSI under conditions of Fe-limitation, was also shown to provide photoprotection to PSII by dissipating excess light energy under oxidative stress conditions (Yeremenko et al., 2004; Ihalainen et al., 2005; Supplementary Table 2).

Given that the ability of cyanobacteria to grow over a large temperature range largely relies on their capacity to optimize the functioning of their photosynthetic apparatus, notably at low temperature that induces a general slowing down of cell metabolism (Murata et al., 2007; Pittera et al., 2014, 2017), we also compared the photophysiology of CRD1 strains and clades I-IV strains at three common growth temperatures. These analyses showed that all three CRD1 strains exhibit a lower growth rate at most temperatures than clade I to IV strains (Fig. 1), possibly explaining why they are easily outcompeted by other taxa when iron is no longer limiting, as observed for instance around the Marquesas Islands (Caputi et al., 2019). Moreover, CRD1 strains also display a lower PSII maximum quantum yield (Fig. 2), suggesting that PSII is partially photoinactivated, that is their D1 repair cycle does not fully compensate damage to this protein, even under optimal growth temperatures. Consistent with this, the very high turnover rate of the D1 protein measured in all CRD1 strains indicates that their PSII is much more sensitive to light stress than other strains and can only trigger a moderate increase in R_PSII_ in response to both light and temperature variations, possibly indicating that they are adapted to live deeper in the water column than clades I to IV. This sensitivity could be partially linked to the abovementioned absence of the OCP, potentially reducing their ability to dissipate excess light energy, although it must be noted that the clade II strain A15-62 also lacks the OCP system. Interestingly in this context, all cold thermotypes including the CRD1B strain BIOS-U3-1, possess more copies of the D1:2 isoform (3-6 copies, average: 3.9 ± 1.1) than warm thermotypes (2-3 copies, average: 2.2 ± 0.4), this isoform providing a lower quantum yield but higher PSII resistance to photoinhibition than D1:1 (Supplementary Table 2; Clarke et al., 1993a, 1993b; Campbell et al., 1995; Garczarek et al., 2008). Moreover, A15-62 is one of the only *Synechococcus* strains to possess two complete copies of the D1:1 isoform, a duplication which could partly explain why, despite the absence of OCP, this atypical clade II strain is able to maintain a high PSII quantum yield over its whole growth temperature range with fairly low D1 repair rates. Interestingly, CRD1 strains also possess a paralog of *psbN*, which was found to be required for the assembly of the PSII reaction centre in *Nicotiana tabacum* and would play an important role in the D1 repair cycle (Torabi et al., 2014).

Taken together, both comparative genomics and photophysiological analyses highlighted a number of specificities of CRD1 strains compared to their clade I-IV counterparts, rather than them possessing traits distinctive of cold or warm thermotypes. In this context, it is worth noting that although strains representative of ESTUs IVA and CRD1B exhibit a similar fundamental thermal niche in culture, the higher median temperature of CRD1B in the field indicates that it preferentially thrives in temperate waters (about 18°C, Fig. 1), where energetically costly temperature adaptation mechanisms might not be essential. This suggests that for members of the CRD1 clade, adaptation to low-Fe conditions likely prevails over adaptation to temperature variations, and/or that adaptation mechanisms to temperature variations might be more complex and diversified than previously thought. Still, in terms of a realized environmental thermal niche, the occurrence of several CRD1 thermotypes likely explains why the CRD1 clade as a whole occupies most Fe-limited areas, a vast ecosystem constituting about 30% of the world Ocean (Moore et al., 2013; Bristow et al., 2017). A notable exception is the Southern Ocean, for which from the little available data shows that *Synechococcus* is scarce south of the polar front (Wilkins et al., 2013; Farrant et al., 2016), consistent with the fairly high low-temperature limit (14°C) of the CRD1B environmental realized niche (Fig. 1), while low-Fe availability likely limits the growth of clades I and IV in this area. In contrast, CRD1 growth does not currently appear to be limited by warm temperatures since most oceanic waters display a temperature below 30°C (Supplementary Fig. 2B; Supplementary Table 1). However, one cannot exclude that with global change, some areas of the world Ocean could become warmer than the highest limits determined here for representative strains of CRD1A and C, i.e. 31°C and 30°C respectively. In this context, it is worth mentioning that in the dataset used for this study, several coastal stations, sampled during the OSD campaign reached 31.5 °C (Supplementary Table 1). Thus, although biogeochemistry global models predict that *Synechococcus* could be one of the winners of the phytoplankton community in a future world Ocean (Flombaum et al., 2013; Schmidt et al., 2020; Visintini et al., 2021), it might well not be able to survive in the warmest low-Fe areas, an ecological niche that is currently expanding (Polovina et al., 2008). Although a few studies have started to analyze the genomic bases of adaptation of *Synechococcus* cells to Fe-limitation in the field (Ahlgren et al., 2020; Garcia et al., 2020), further comparative genomic and physiological studies are still needed to decipher the specific capacity of CRD1 clade members to deal with Fe-limitation which should help predict the future distribution and dynamics of *Synechococcus* taxa in the world Ocean.

## DATA AVAILABILITY STATEMENT

Unpublished metabarcoding data supporting the conclusions of this article are available as raw data (SRA accession numbers) and processed data (number of *petB* reads per ESTU) in Supplementary Table 1. The latter Table also encompasses the description of all environmental samples used in this study.

## AUTHOR CONTRIBUTIONS

MF, HD and LG designed the experiments. MF, LD, HD, MR, AG, FRJ, TS and GM collected the samples and performed the physiological measurements. MF and DM ran the flow cytometry analyses. FL isolated several CRD1 strains used in this study. HD, XX, DJS, HL and LG performed sequencing and bioinformatics analyses of metabarcodes. MH, EC, FP and LG developed and refined the Cyanorak *v2.1* database. MF, HD, FP and LG made the figures. MF, LD, HD, SB, FP and LG interpreted results. All the authors contributed to the preparation of the manuscript, read and approved the final manuscript.

## ACKNOWLEDGEMENTS

We would like to thank Thierry Cariou for providing physico-chemical parameters from the SOMLIT-Astan station and Gwenn Tanguy (Biogenouest genomics core facility) and Monica Moniz for their sequencing of *petB* metabarcodes. We are also most grateful to Nathalie Simon for coordinating the phytoplankton time-series at the SOMLIT-Astan station, Christophe Six for technical hints on PAM fluorimetry and HPLC analyses as well as Priscillia Gourvil and Martin Gachenot from the Roscoff Culture Collection (http://roscoff-culture-collection.org/) and Florian Humily for isolating and/or maintaining the *Synechococcus* strains used in this study. We also thank the support and commitment of the *Tara* Oceans coordinators and consortium, Agnès b. and E. Bourgois, the Veolia Environment Foundation, Région Bretagne, Lorient Agglomeration, World Courier, Illumina, the EDF Foundation, FRB, the Prince Albert II de Monaco Foundation, the *Tara* schooner and its captains and crew.

## SUPPLEMENTARY MATERIAL

**Supplementary Table 1** | Environmental samples used in this study for the determination of the realized environmental niches of the major *Synechococcus* ESTUs

**Supplementary Table 2** | Phyletic pattern of CRD1 and clade I-IV genes mentioned in this study retrieved from the Cyanorak v2.1 database

## REFERENCES

Ahlgren, N. A., Belisle, B. S., and Lee, M. D. (2020). Genomic mosaicism underlies the adaptation of marine *Synechococcus* ecotypes to distinct oceanic iron niches. Environ. Microbiol. 22, 1801–1815. doi:10.1111/1462-2920.14893.

Ahlgren, N. A., and Rocap, G. (2012). Diversity and distribution of marine *Synechococcus:* Multiple gene phylogenies for consensus classification and development of qPCR assays for sensitive measurement of clades in the ocean. Front. Microbiol. 3, 213–213. doi:10.3389/fmicb.2012.00213.

Biller, S. J., Berube, P. M., Lindell, D., and Chisholm, S. W. (2015). *Prochlorococcus:* The structure and function of collective diversity. Nat. Rev. Microbiol. 13, 13–27. doi:10.1038/nrmicro3378.

Breton, S., Jouhet, J., Guyet, U., Gros, V., Pittera, J., Demory, D., et al. (2020). Unveiling membrane thermoregulation strategies in marine picocyanobacteria. New Phytol. 225, 2396–2410. doi:10.1111/nph.16239.

Bristow, L. A., Mohr, W., Ahmerkamp, S., and Kuypers, M. M. M. (2017). Nutrients that limit growth in the ocean. Curr. Biol. 27, R431–R510. doi:10.1016/j.cub.2017.03.030.

Campbell, D., Hurry, V., Clarke, A. K., Gustafsson, P., and Öquist, G. (1998). Chlorophyll fluorescence analysis of cyanobacterial photosynthesis and acclimation. Microbiol. Mol. Biol. Rev. 62, 667–683. doi:10.1128/mmbr.62.3.667-683.1998.

Campbell, D., Zhou, G., Gustafsson, P., Oquist, G., and Clarke, A. K. (1995). Electron transport regulates exchange of two forms of photosystem II D1 protein in the cyanobacterium *Synechococcus*. EMBO J. 14, 5457–5466. doi:10.1128/MMBR.62.3.667-683.1998.

Caputi, L., Carradec, Q., Eveillard, D., Kirilovsky, A., Pelletier, E., Pierella Karlusich, J. J., et al. (2019). Community-level responses to iron availability in open ocean plankton ecosystems. Glob. Biogeochem. Cycles 33, 391–419. doi:10.1029/2018GB006022.

Clarke, A. K., Hurry, V. M., Gustafsson, P., and Oquist, G. (1993a). Two functionally distinct forms of the photosystem II reaction-center protein D1 in the cyanobacterium *Synechococcus* sp. PCC 7942. Proc. Natl. Acad. Sci. U. S. A. 90, 11985–11989. doi:10.1073/pnas.90.24.11985.

Clarke, A. K., Soitamo, A., Gustafsson, P., and Oquist, G. (1993b). Rapid interchange between two distinct forms of cyanobacterial photosystem II reaction-center protein D1 in response to photoinhibition. Proc. Natl. Acad. Sci. U. S. A. 90, 9973–9977. doi:10.1073/pnas.90.21.9973.

Claustre, H., Sciandra, A., and Vaulot, D. (2008). Introduction to the special section bio-optical and biogeochemical conditions in the South East Pacific in late 2004: the BIOSOPE program. Biogeosciences 5, 679–691. doi:10.5194/bg-5-679-2008.

Delmont, T. O., Gaia, M., Hinsinger, D. D., Fremont, P., Guerra, A. F., Eren, A. M., et al. (2020). Functional repertoire convergence of distantly related eukaryotic plankton lineages revealed by genome-resolved metagenomics. BioRxiv, 2020.10.15.341214-2020.10.15.341214. doi:10.1101/2020.10.15.341214.

Doré, H., Farrant, G. K., Guyet, U., Haguait, J., Humily, F., Ratin, M., et al. (2020). Evolutionary mechanisms of long-term genome diversification associated with niche partitioning in marine picocyanobacteria. Front. Microbiol. 11, 567431. doi:10.3389/fmicb.2020.567431.

Doré, H., Leconte, Jade, Breton, Solène, Demory, David, Hoebeke, Mark, Corre, Erwan, et al. (2022). Global phylogeography of marine *Synechococcus* in coastal areas unveils strikingly different communities than in open ocean. BioRxiv.

Dufresne, A., Ostrowski, M., Scanlan, D. J., Garczarek, L., Mazard, S., Palenik, B. P., et al. (2008). Unraveling the genomic mosaic of a ubiquitous genus of marine cyanobacteria. Genome Biol. 9, R90. doi:10.1186/gb-2008-9-5-r90.

Farrant, G. K., Doré, H., Cornejo-Castillo, F. M., Partensky, F., Ratin, M., Ostrowski, M., et al. (2016). Delineating ecologically significant taxonomic units from global patterns of marine picocyanobacteria. Proc. Natl. Acad. Sci. U. S. A. 113, E3365–E3374. doi:10.1073/pnas.1524865113.

Findlay, V. J., Tapiero, H., and Townsend, D. M. (2005). Sulfiredoxin: a potential therapeutic agent? Biomed. Pharmacother. 59, 374–379. doi:10.1016/j.biopha.2005.07.003.

Flombaum, P., Gallegos, J. L., Gordillo, R. a, Rincón, J., Zabala, L. L., Jiao, N., et al. (2013). Present and future global distributions of the marine Cyanobacteria *Prochlorococcus* and *Synechococcus*. Proc. Natl. Acad. Sci. U. S. A. 110, 9824–9829. doi:10.1073/pnas.1307701110.

Garcia, C. A., Hagstrom, G. I., Larkin, A. A., Ustick, L. J., Levin, S. A., Lomas, M. W., et al. (2020). Linking regional shifts in microbial genome adaptation with surface ocean biogeochemistry. Philos. Trans. R. Soc. B Biol. Sci. 375, 20190254. doi:10.1098/rstb.2019.0254.

Garczarek, L., Dufresne, A., Blot, N., Cockshutt, A. M., Peyrat, A., Campbell, D. A., et al. (2008). Function and evolution of the *psbA* gene family in marine *Synechococcus*: *Synechococcus* sp. WH7803 as a case study. ISME J. 2, 937–953. doi:10.1038/ismej.2008.46.

Garczarek, L., Guyet, U., Doré, H., Farrant, G. K., Hoebeke, M., Brillet-Guéguen, L., et al. (2021). Cyanorak v2.1: a scalable information system dedicated to the visualization and expert curation of marine and brackish picocyanobacteria genomes. Nucleic Acids Res. 49, D667–D676. doi:10.1093/nar/gkaa958.

Gutiérrez-Rodríguez, A., Slack, G., Daniels, E. F., Selph, K. E., Palenik, B., and Landry, M. R. (2014). Fine spatial structure of genetically distinct picocyanobacterial populations across environmental gradients in the Costa Rica Dome. Limnol. Oceanogr. 59, 705–723. doi:10.4319/lo.2014.59.3.0705.

Guyet, U., Nguyen, N. A., Doré, H., Haguait, J., Pittera, J., Conan, M., et al. (2020). Synergic effects of temperature and irradiance on the physiology of the marine *Synechococcus* strain WH7803. Front. Microbiol. 11, 1707. doi:10.3389/fmicb.2020.01707.

Humily, F., Partensky, F., Six, C., Farrant, G. K., Ratin, M., Marie, D., et al. (2013). A gene island with two possible configurations is involved in chromatic acclimation in marine *Synechococcus*. PLoS One 8, e84459. doi:10.1371/journal.pone.0084459.

Ihalainen, J. A., D’Haene, S., Yeremenko, N., van Roon, H., Arteni, A. A., Boekema, E. J., et al. (2005). Aggregates of the chlorophyll-binding protein IsiA (CP43’) dissipate energy in cyanobacteria. Biochemistry 44, 10846–10853. doi:10.1021/bi0510680.

Inoue, N., Taira, Y., Emi, T., Yamane, Y., Kashino, Y., Koike, H., et al. (2001). Acclimation to the growth temperature and the high-temperature effects on photosystem II and plasma membranes in a mesophilic cyanobacterium *Synechocystis* sp. PCC6803. Plant Cell Physiol. 42, 1140–1148. doi:10.1093/pcp/pce147.

Johnson, Z. I., Zinser, E. R., Coe, A., McNulty, N. P., Woodward, E. M. S., and Chisholm, S. W. (2006). Niche partitioning among *Prochlorococcus* ecotypes along ocean-scale environmental gradients. Science 311, 1737–1740. doi:10.1126/science.1118052.

Kana, T. M., Glibert, P. M., Goericke, R., and Welschmeyer, N. A. (1988). Zeaxanthin and ß-carotene in *Synechococcus* WH7803 respond differently to irradiance. Limnol. Oceanogr. 33, 1623–1626. doi:10.4319/lo.1988.33.6part2.1623.

Kashtan, N., Roggensack, S. E., Rodrigue, S., Thompson, J. W., Biller, S. J., Coe, A., et al. (2014). Single-cell genomics reveals hundreds of coexisting subpopulations in wild *Prochlorococcus*. Science 344, 416–420. doi:10.1126/science.1248575.

Kent, A. G., Baer, S. E., Mouginot, C., Huang, J. S., Larkin, A. A., Lomas, M. W., et al. (2019). Parallel phylogeography of *Prochlorococcus* and *Synechococcus*. ISME J. 13, 430–441. doi:10.1038/s41396-018-0287-6.

Kettler, G. C., Martiny, A. C., Huang, K., Zucker, J., Coleman, M. L., Rodrigue, S., et al. (2007). Patterns and implications of gene gain and loss in the evolution of *Prochlorococcus*. PLoS Genet. 3, e231. doi:10.1371/journal.pgen.0030231.

Kirilovsky, D. (2007). Photoprotection in cyanobacteria: the orange carotenoid protein (OCP)-related non-photochemical-quenching mechanism. Photosynth. Res. 93, 7. doi:10.1007/s11120-007-9168-y.

Larkin, A. A., Blinebry, S. K., Howes, C., Lin, Y., Loftus, S. E., Schmaus, C. A., et al. (2016). Niche partitioning and biogeography of high light adapted *Prochlorococcus* across taxonomic ranks in the North Pacific. ISME J. 10, 1555–1567. doi:10.1038/ismej.2015.244.

Larkin, A. A., and Martiny, A. C. (2017). Microdiversity shapes the traits, niche space, and biogeography of microbial taxa: The ecological function of microdiversity. Environ. Microbiol. Rep. 9, 55–70. doi:10.1111/1758-2229.12523.

Latifi, A., Ruiz, M., and Zhang, C. C. (2009). Oxidative stress in cyanobacteria. FEMS Microbiol. Rev. 33, 258-278–258–278. doi:10.1111/j.1574-6976.2008.00134.x.

Mackey, K. R. M., Paytan, A., Caldeira, K., Grossman, A. R., Moran, D., McIlvin, M., et al. (2013). Effect of temperature on photosynthesis and growth in marine *Synechococcus* spp. Plant Physiol. 163, 815–829. doi:10.1104/pp.113.221937.

Marie, D., Partensky, F., Vaulot, D., and Brussaard, C. (1999). Enumeration of phytoplankton, bacteria, and viruses in marine samples. Curr. Protoc. Cytom. 10, 11.11.1–11.11.15. doi:10.1002/0471142956.cy1111s10.

Mella-Flores, D., Mazard, S., Humily, F., Partensky, F., Mahé, F., Bariat, L., et al. (2011). Is the distribution of *Prochlorococcus* and *Synechococcus* ecotypes in the Mediterranean Sea affected by global warming? Biogeosciences 8, 2785–2804. doi:10.5194/bg-8-2785-2011.

Mikami, K., and Murata, N. (2003). Membrane fluidity and the perception of environmental signals in cyanobacteria and plants. Prog. Lipid Res. 42, 527–543. doi:10.1016/S0163-7827(03)00036-5.

Moore, C. M., Mills, M. M., Arrigo, K. R., Berman-Frank, I., Bopp, L., Boyd, P. W., et al. (2013). Processes and patterns of oceanic nutrient limitation. Nat. Geosci. 6, 701–710. doi:10.1038/ngeo1765.

Murata, N., Takahashi, S., Nishiyama, Y., and Allakhverdiev, S. I. (2007). Photoinhibition of photosystem II under environmental stress. Biochim. Biophys. Acta - Bioenerg. 1767, 414–421. doi:10.1016/j.bbabio.2006.11.019.

Nishiyama, Y., Allakhverdiev, S. I., and Murata, N. (2006). A new paradigm for the action of reactive oxygen species in the photoinhibition of photosystem II. Biochim. Biophys. Acta BBA - Bioenerg. 1757, 742–749. doi:10.1016/j.bbabio.2006.05.013.

Paulsen, M. L., Doré, H., Garczarek, L., Seuthe, L., Müller, O., Sandaa, R.-A., et al. (2016). *Synechococcus* in the Atlantic gateway to the Arctic Ocean. Front. Mar. Sci. 3, 191. doi:10.3389/fmars.2016.00191.

Pearman, P. B., Guisan, A., Broennimann, O., and Randin, C. F. (2008). Niche dynamics in space and time. Trends Ecol. Evol. 23, 149–158. doi:10.1016/j.tree.2007.11.005.

Pittera, J., Humily, F., Thorel, M., Grulois, D., Garczarek, L., and Six, C. (2014). Connecting thermal physiology and latitudinal niche partitioning in marine *Synechococcus*. ISME J. 8, 1221–1236. doi:10.1038/ismej.2013.228.

Pittera, J., Jouhet, J., Breton, S., Garczarek, L., Partensky, F., Maréchal, É., et al. (2018). Thermoacclimation and genome adaptation of the membrane lipidome in marine *Synechococcus*. Environ. Microbiol. 20, 612–631. doi:10.1111/1462-2920.13985.

Pittera, J., Partensky, F., and Six, C. (2017). Adaptive thermostability of light-harvesting complexes in marine picocyanobacteria. ISME J. 11, 112–124. doi:10.1038/ismej.2016.102.

Polovina, J. J., Howell, E. A., and Abecassis, M. (2008). Ocean’s least productive waters are expanding. Geophys. Res. Lett. 35, L03618. doi:10.1029/2007GL031745.

Rippka, R., Coursin, T., Hess, W., Lichtle, C., Scanlan, D. J., Palinska, K. A., et al. (2000). *Prochlorococcus marinus* Chisholm et al. 1992 subsp. *pastoris* subsp. nov. strain PCC 9511, the first axenic chlorophyll *a*2/*b*2-containing cyanobacterium (Oxyphotobacteria). Int. J. Syst. Evol. Microbiol. 50, 1833–1847. doi:10.1099/00207713-50-5-1833.

Saito, M. A., Rocap, G., and Moffett, J. W. (2005). Production of cobalt binding ligands in a *Synechococcus* feature at the Costa Rica upwelling dome. Limnol. Oceanogr. 50, 279–290. doi:10.4319/lo.2005.50.1.0279.

Scanlan, D. J., Ostrowski, M., Mazard, S., Dufresne, A., Garczarek, L., Hess, W. R., et al. (2009). Ecological genomics of marine picocyanobacteria. Microbiol. Mol. Biol. Rev. 73, 249–299. doi:10.1128/MMBR.00035-08.

Schloss, P. D., Westcott, S. L., Ryabin, T., Hall, J. R., Hartmann, M., Hollister, E. B., et al. (2009). Introducing Mothur: Open-source, platform-independent, community-supported software for describing and comparing microbial communities. Appl. Environ. Microbiol. 75, 7537–7541. doi:10.1128/AEM.01541-09.

Schmidt, K., Birchill, A. J., Atkinson, A., Brewin, R. J. W., Clark, J. R., Hickman, A. E., et al. (2020). Increasing picocyanobacteria success in shelf waters contributes to long-term food web degradation. Glob. Change Biol. 26, 5574–5587. doi:10.1111/gcb.15161.

Six, C., Joubin, L., Partensky, F., Holtzendorff, J., and Garczarek, L. (2007). UV-induced phycobilisome dismantling in the marine picocyanobacterium *Synechococcus* sp. WH8102. Photosynth. Res. 92, 75–86. doi:10.1007/s11120-007-9170-4.

Six, C., Ratin, M., Marie, D., and Corre, E. (2021). Marine *Synechococcus* picocyanobacteria: Light utilization across latitudes. Proc. Natl. Acad. Sci. U. S. A. 118, e2111300118. doi:10.1073/pnas.2111300118.

Six, C., Thomas, J., Brahamsha, B., Lemoine, Y., and Partensky, F. (2004). Photophysiology of the marine cyanobacterium *Synechococcus* sp. WH8102, a new model organism. Aquat. Microb. Ecol. 35, 17–29. doi:10.3354/ame035017.

Six, C., Thomas, J.-C., Thion, L., Lemoine, Y., Zal, F., and Partensky, F. (2005). Two novel phycoerythrin-associated linker proteins in the marine cyanobacterium *Synechococcus* sp. strain WH8102. J. Bacteriol. 187, 1685–1694. doi:10.1128/JB.187.5.1685-1694.2005.

Sohm, J. A., Ahlgren, N. A., Thomson, Z. J., Williams, C., Moffett, J. W., Saito, M. A., et al. (2016). Co-occurring *Synechococcus* ecotypes occupy four major oceanic regimes defined by temperature, macronutrients and iron. ISME J. 10, 333–345. doi:10.1038/ismej.2015.115.

Sunagawa, S., Coelho, L. P., Chaffron, S., Kultima, J. R., Labadie, K., Salazar, G., et al. (2015). Structure and function of the global ocean microbiome. Science 348, 1261359–1261359. doi:10.1126/science.1261359.

Torabi, S., Umate, P., Manavski, N., Plöchinger, M., Kleinknecht, L., Bogireddi, H., et al. (2014). PsbN is required for assembly of the photosystem II reaction center in *Nicotiana tabacum*. Plant Cell 26, 1183–1199. doi:10.1105/tpc.113.120444.

Umena, Y., Kawakami, K., Shen, J.-R., and Kamiya, N. (2011). Crystal structure of oxygen-evolving photosystem II at a resolution of 1.9 Å. Nature 473, 55–60. doi:10.1038/nature09913.

Visintini, N., Martiny, A. C., and Flombaum, P. (2021). *Prochlorococcus, Synechococcus*, and picoeukaryotic phytoplankton abundances in the global ocean. Limnol. Oceanogr. Lett. 6, 207–215. doi:10.1002/lol2.10188.

Wilkins, D., Lauro, F. M., Williams, T. J., Demaere, M. Z., Brown, M. V., Hoffman, J. M., et al. (2013). Biogeographic partitioning of Southern Ocean microorganisms revealed by metagenomics. Environ. Microbiol. 15, 1318–1333. doi:10.1111/1462-2920.12035.

Xia, X., Cheung, S., Endo, H., Suzuki, K., and Liu, H. (2019). Latitudinal and vertical variation of *Synechococcus* assemblage composition along 170°W transect from the South Pacific to the Arctic Ocean. Microb. Ecol. 77, 333–342. doi:10.1007/s00248-018-1308-8.

Xu, W., and Wang, Y. (2017). “Function and structure of cyanobacterial photosystem I,” in Photosynthesis: Structures, Mechanisms, and Applications, eds. H. J. M. Hou, M. M. Najafpour, G. F. Moore, and S. I. Allakhverdiev (Cham: Springer International Publishing), 111–168. doi:10.1007/978-3-319-48873-8_7.

Yeremenko, N., Kouřil, R., Ihalainen, J. A., D’Haene, S., van Oosterwijk, N., Andrizhiyevskaya, E. G., et al. (2004). Supramolecular organization and dual function of the IsiA chlorophyll-binding protein in cyanobacteria. Biochemistry 43, 10308–10313. doi:10.1021/bi048772l.

Zwirglmaier, K., Jardillier, L., Ostrowski, M., Mazard, S., Garczarek, L., Vaulot, D., et al. (2008). Global phylogeography of marine *Synechococcus* and *Prochlorococcus* reveals a distinct partitioning of lineages among oceanic biomes. Environ. Microbiol. 10, 147–161. doi:https://doi.org/10.1111/j.1462-2920.2007.01440.x.

