## Supplemental Figures for "Comparative thermophysiology of marine *Synechococcus* CRD1 strains isolated from different thermal niches in iron-depleted areas"

### Ferrieux et al.

### Supplementary Figures

frontiers

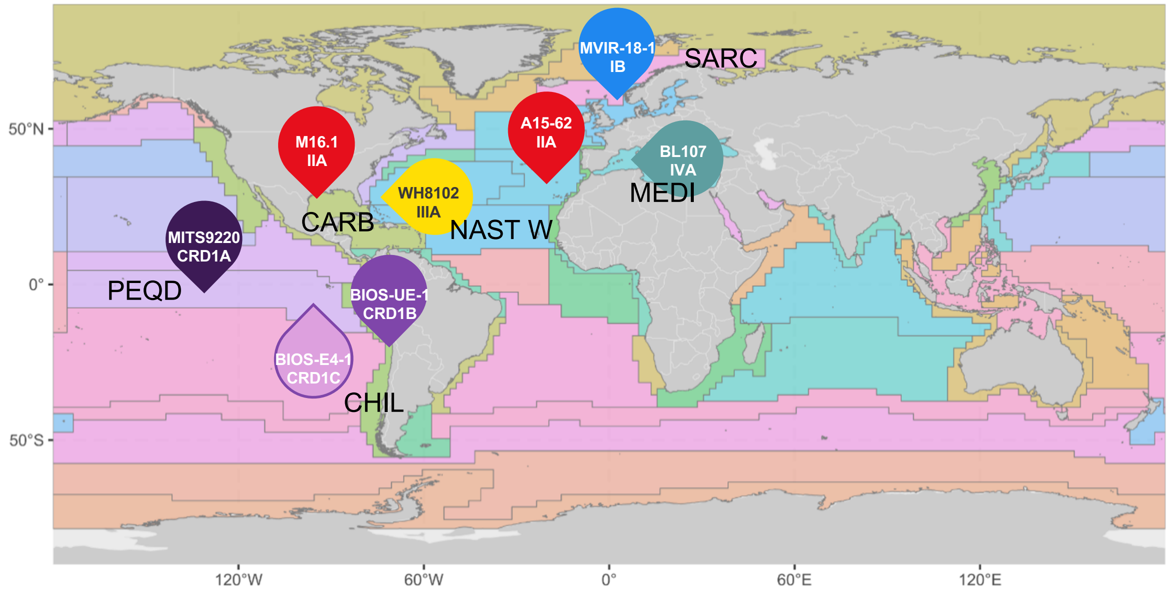


**Supplementary Figure 1. Isolation sites of the *Synechococcus* strains used in this study.** Isolation site of each strain is indicated on the map by a bubble arrow colored according to their corresponding ESTUs indicated below the strain name. Longhurst provinces (Longhurst A., 2007, Ecological Geography of the Sea, Academic Press, London) are shown as a colored background shown in the insert. Only provinces from which at least one strain has been isolated are indicated on the map using the following abbreviations: PEQD (Pacific equatorial divergence, Pacific, Trade wind), CHIL (Chile-Peru, current coastal province, Pacific, Coastal), NAST W (Northwest Atlantic subtropical gyral, Atlantic, Westerly), CARB (Caribbean, Atlantic, Trade wind), MEDI (Mediterranean Sea, Atlantic, Westerly), SARC (Atlantic sub-Arctic, Atlantic, Polar).

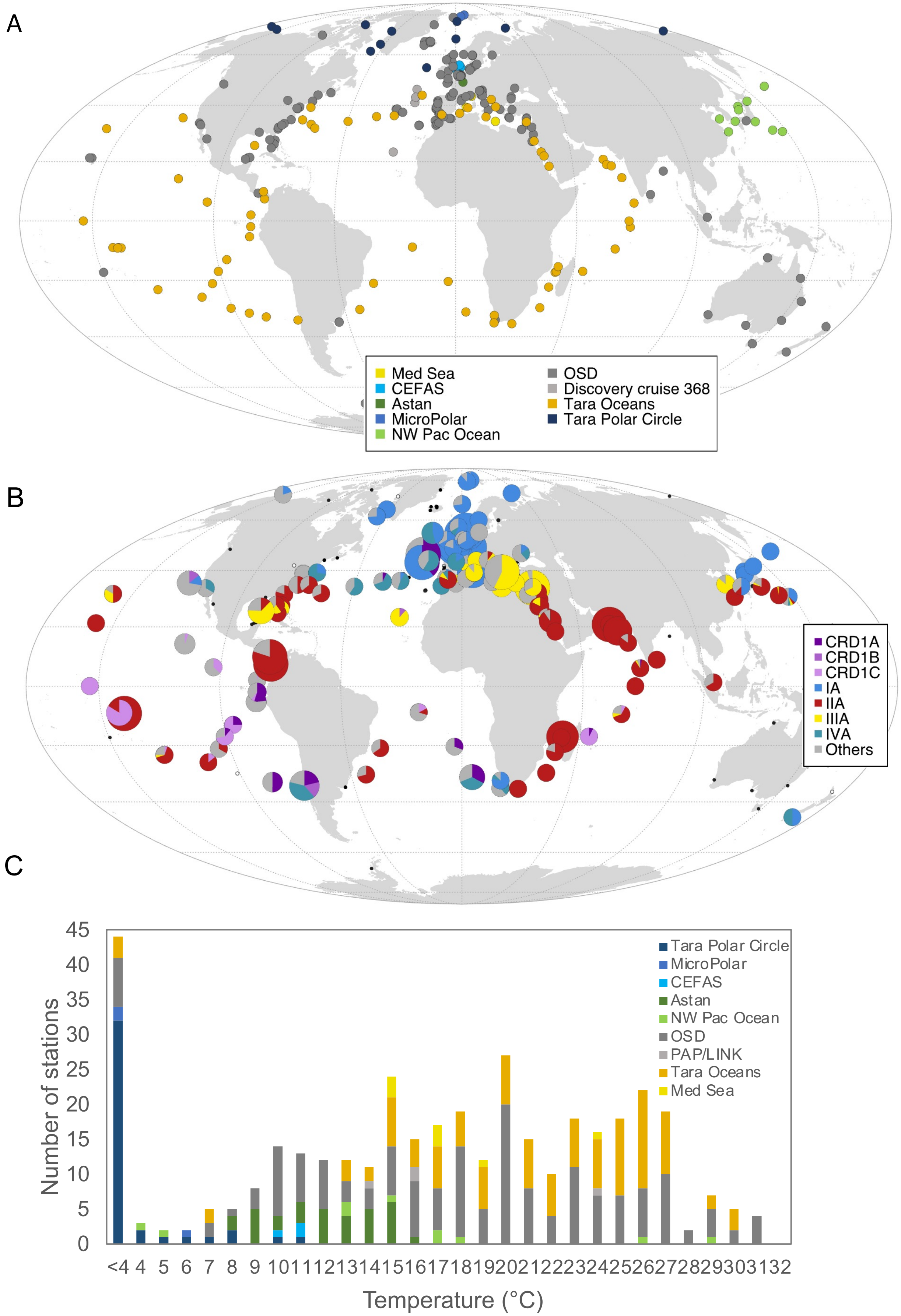


**Supplementary Figure 2. Oceanwide environmental data used in this study to determine the environmental realized thermal niches of the main ESTUs from clades CRD1 and I to IV. (A)** Map of the sampling sites, **(B)** Relative abundance of *Synechococcus* ESTUs IA to IVA and CRD1A to C. **(C)** Temperature distribution of the sampling sites. The inserts specifies **(A,C)** the name of campaigns or datasets analyzed and **(B)** the *Synechococcus* ESTUs.

A. RpcA

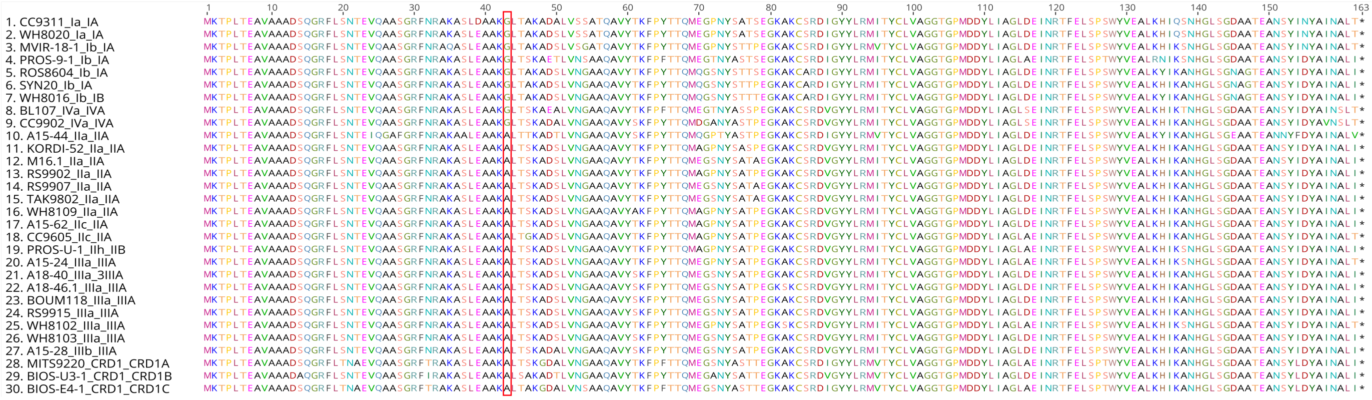


B. RpcB

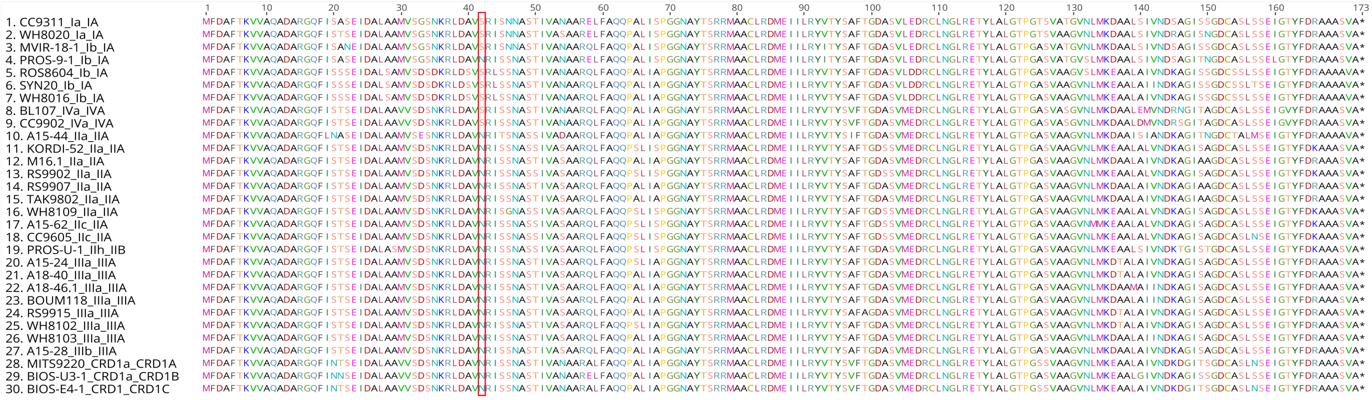


**Supplementary Figure 3. Alignment of RpcA and RpcB, encoding phycocyanin α- and β-subunits, from CRD1 and clades I-IV *Synechococcus* strains. (A)** RpcA. **(B)** RpcB. Substitutions potentially involved in thermotolerance are shown by a red rectangle in the alignment.

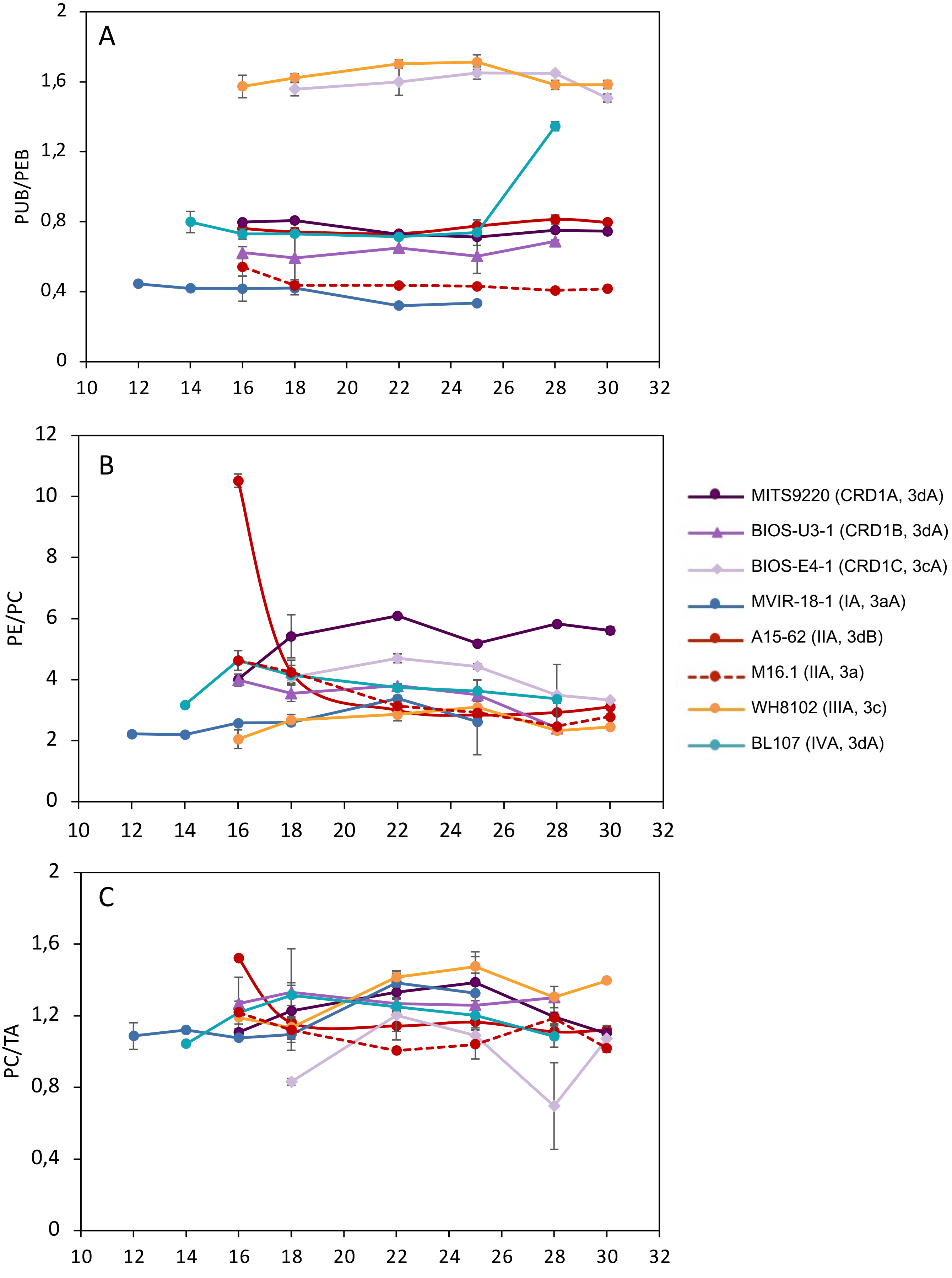


**Supplementary Figure 4. Variation with growth temperature of phycobilins and phycobiliproteins fluorescence excitation and emission ratios. (A)** Average PUB:PEB ratios. **(B)** Average phycoerythrin (PE) to phycocyanin (PC) ratios. **(C)** Average PC to terminal acceptor (TA) ratios. The insert indicates the strain names, their corresponding ESTU (*sensu* Farrant et al., 2016) and pigment type (*sensu* Humily et al., 2013) between brackets.

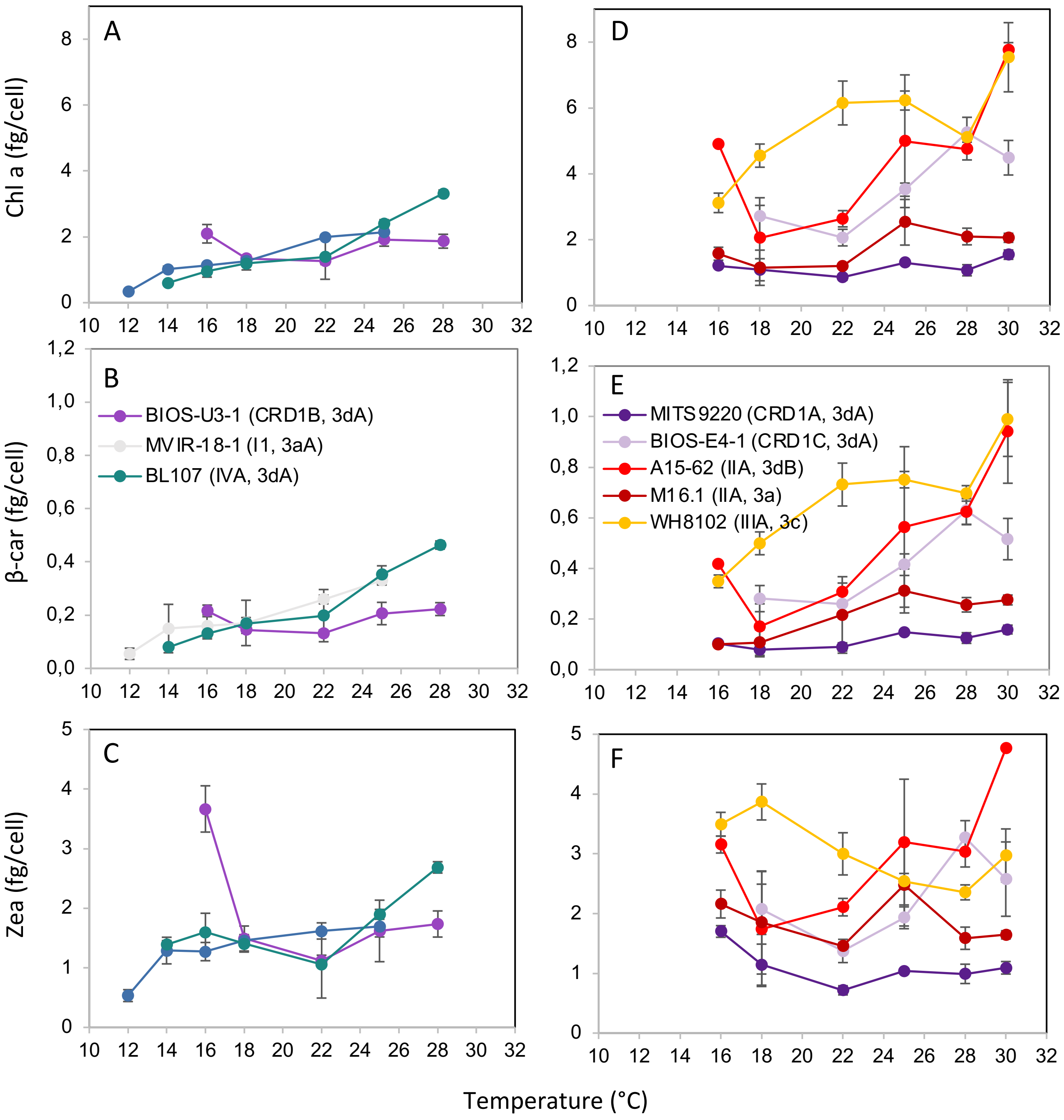


**Supplementary Figure 5. Variation with growth temperature of the three main liposoluble pigments per cell for CRD1 vs. clade I and IV strains. (A, D)** Chlorophyll (Chl) *a* content (fg/cell). **(B, E)** Zeaxanthin (Zea) content (fg/cell). **(C, F)** β-carotene (β-car) content (fg/cell). **(A-C)** CRD1-B strain BIOS-U3-1 *vs.* cold thermotypes. **(D-F)** CRD1-A strain MITS9220 and CRD1-C strain BIOS-E4-1 *vs.* warm thermotypes. Inserts indicate the strain names and their corresponding ESTU (*sensu* Farrant et al., 2016) between brackets.

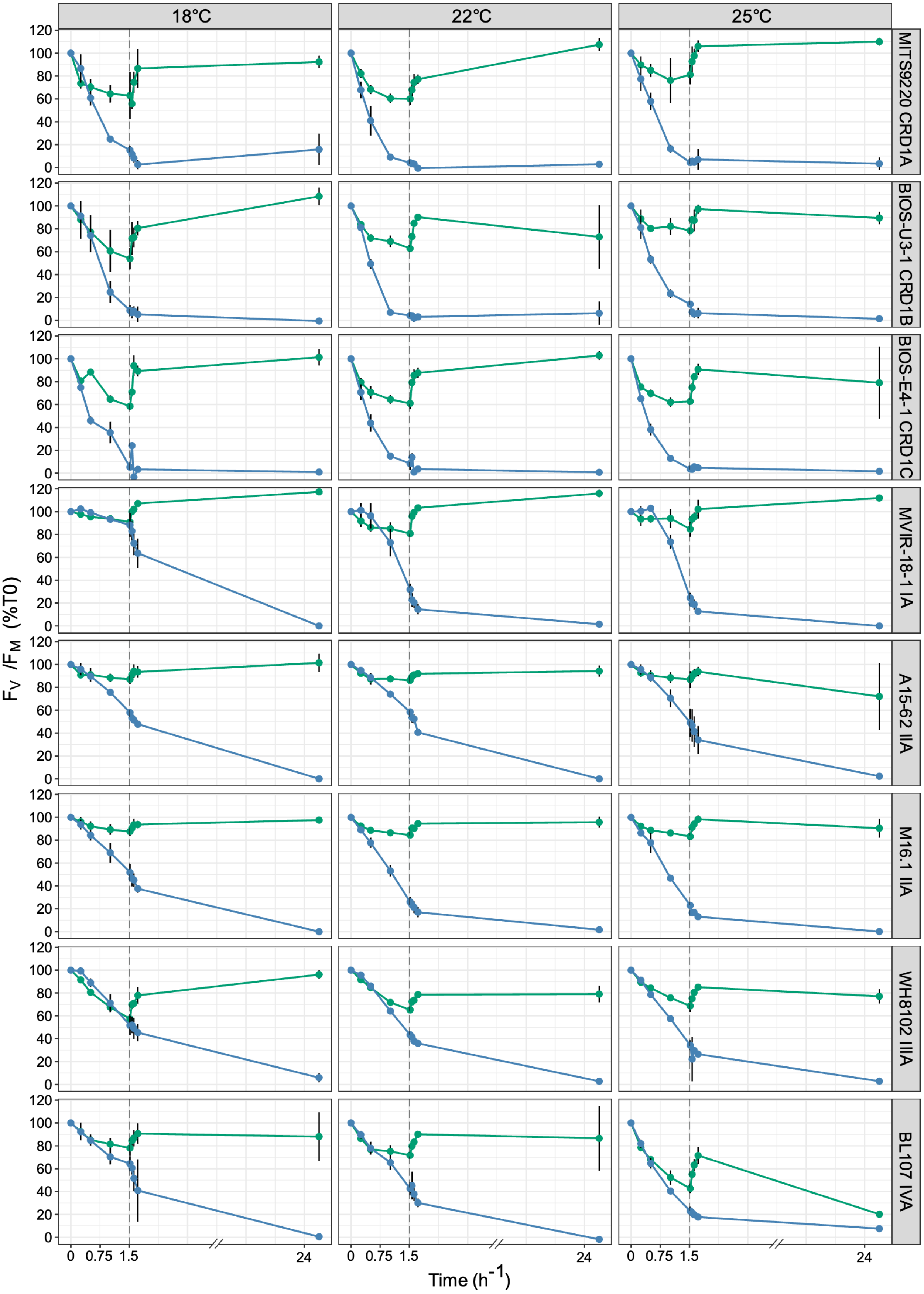


**Supplementary Figure 6. Time course of photosystem II quantum yield ($F_V/F_M$) following light stress in the presence or absence of lincomycin for CRD1 and clade I-IV strains acclimated to different temperatures.** Cultures acclimated to 75 µE $m^{-2}$ $s^{-1}$ were shifted to 375 µE $m^{-2}$ $s^{-1}$ at T0 for 90 min, then shifted back to the initial light conditions for 24h as indicated by a vertical dashed line on each figure. Strain names and their corresponding ESTU between brackets (*sensu* Farrant et al., 2016) are indicated on the right hand side, acclimation temperature are indicated on the top, whilst line colour indicates the lincomycin treatment (i.e. -/+ linco).
